# Exhaled breath profiling for non-invasive monitoring of cognitive functioning in children

**DOI:** 10.1101/2025.03.25.645320

**Authors:** Joris Meurs, Ben Henderson, Claudia van Dun, Guilherme Lopes Batitsta, Evangelia Sakkoula, Janna A. van Diepen, Gabriele Gross, Esther Aarts, Simona M. Cristescu

## Abstract

**Introduction:** Childhood is a critical period for the development of executive functioning skills, including selective attention and inhibitory control, which are essential for cognitive development. Optimal brain development during this time requires appropriate levels of macronutrient intake. Metabolomics can offer valuable insights into which metabolites cognitive functioning and the underlying gut-brain interactions.

**Objectives:** This study aimed to explore to use of breathomics to investigate associations between exhaled metabolites and executive functioning in children.

**Methods:** Children (8-10 years; N = 31) were recruited via flyers at schools and after-school care. The assessment of executive functioning was done using Eriksen flanker task. Breath samples were collected in Tedlar® bags and analyzed with proton transfer reaction–mass spectrometry (PTR-MS). On-breath peaks were selected and subjected to partial least squares (PLS) regression. Significance multivariate correlation (sMC) was used afterwards to select metabolites bearing predictive power towards executive functioning.

**Results:** Gut microbiome-related metabolites (methane, ethanol, and butyric acid) present in exhaled breath were associated with an improved executive functioning, whereas isoprene was linked to reduced executive functioning. Additionally, increased levels of inflammatory markers, ethylene and acetaldehyde, were associated with a higher compatibility effect in error rates, suggesting diminished cognitive control. These VOCs were putatively linked with specific gut microbial taxa; for instance, reduced *Bacteroidetes* abundance (associated with methane production) is associated with decreased inhibitory control, while *Enterobacteriaceae* were linked to lipopolysaccharide-induced inflammation which is also a process that causes increased ethylene production.

**Conclusion:** This proof-of-concept study demonstrates that VOCs in exhaled breath could serve as a promising non-invasive tool for assessing gut-brain interactions related to executive functioning in children.

## Introduction

The development of executive functioning skills, such as selective attention and inhibitory control, is important during childhood as these skills are essential for cognitive development (White and Greenfield 2017). Optimal brain development during this period requires adequate levels of macronutrients, which can have immediate effects on cognitive performance (Hoyland et al. 2008; Leigh Gibson and Green 2002; Muth and Park 2021). The gut-brain axis (GBA) may plan a significant role in this process, linking the gut microbiota with the central nervous system (CNS) (Zhang et al. 2020). The gut microbiota are important for digestive function, immune response, and metabolism, and more recently have been suggested to also play a role in neuropsychiatric health (Cryan et al. 2020). Diet significantly influences this relationship, with certain foods promoting specific taxa within the gut microbial community, which in turn can affect the CNS (Kim 2024). There is growing evidence suggesting an association between the GBA and children’s cognition (Naspolini et al. 2024; S. M.-S. Tran and Mohajeri 2021). with certain microbial metabolites, like short-chain fatty acids (SCFAs) being important mediators in gut-brain communication (Dalile et al. 2019; Majumdar et al. 2023; O’Riordan et al. 2022).

Traditionally, monitoring of metabolites produced by the gut microbiota, such as SCFAs, is done in blood or faecal samples. However, blood sampling is an invasive procedure and faecal concentrations may not accurately reflect *in vivo* concentrations and/or production rates (den Besten et al. 2013). As an alternative, volatile organic compounds (VOCs) in exhaled breath offer a non-invasive method to gain insights into various metabolic processes (Henderson et al. 2022). Exhaled breath is readily available and provides a snapshot of the body’s metabolic state (Buszewski et al. 2007). Previous studies have found strong (in)direct relations between exhaled VOCs and the gut microbiome such as in gastrointestinal disease and nutritional status (Bhandari et al. 2023; Rondanelli et al. 2019; Smolinska et al. 2018). VOCs produced by the gut microbiome can directly or indirectly, through VOC emission from faeces, enter the bloodstream and can therefore also be observed in exhaled breath (Rondanelli et al. 2019), highlighting the potential of exhaled breath analyses to explore gut-brain communication and cognitive functioning in children.

Given that the developing brain is particularly sensitive to the effects of diet on cognitive functioning (Murray and Chen 2019), it is yet not fully understood how diet affects cognition during childhood. Executive functioning, which includes the ability to plan, focus, memorise instructions and deal with multiple tasks (Kavanaugh et al. 2019) is a critical aspect of cognition. The Eriksen flanker task is a widely used tool to assess executive functioning(Eriksen and Eriksen 1974), measuring selective attention and cognitive/inhibitory control by comparing reaction times and accuracy between congruent and incongruent trials.

This study explored the analysis of exhaled breath as a non-invasive tool to identify associations between VOCs produced by the gut microbiome and executive cognitive functioning. This cross-sectional study aims to provide preliminary insights into how exhaled breath analysis might provide mechanistic insights into effects of the gut microbiota on executive functioning and its potential for future studies on gut-brain communication.

## 2. Materials & Methods

### 2.1 Study design

The study has a cross-sectional design targeting healthy Dutch primary school children. Approval of the study design was given by the medical ethical committee of the Radboudumc and registered under NL64464.091.17. The study complied with the Declaration of Helsinki. Children between 8 and 10 years old (*N* = 31) were recruited via flyers at schools and after-school care. We carried out both over-the-phone and in-person screening to ensure children fit the inclusion criteria having no: sensory-motor handicaps, psychiatric disorders, diabetes, chronic inflammation disease, daily medication (including ibuprofen and aspirin, but excluding homeopathic and vitamin supplements), tooth extractions in the past month, yeast infection in the mouth, antibiotics use in the past three months, drastic changes in diet in the past year, unremovable metal in the body, active implant, pacemaker, neurostimulator, insulin pump or hearing device, epilepsy, claustrophobia, history of brain surgery; unremovable dental wire, vaccination in the past month. Testing sessions took place in the afternoon, between 13.00h and 17.30h. Children that did not complete the procedure (i.e. absence of breath sample or flanker task results) were excluded for further analysis. This resulted in a total of 26 participants. Demographic data for the included study participants is summarised in Table 1. Parents of the participants received a 25-minute online Food Frequency Questionnaire (FFQ) assessing the child’s daily habitual food intake in the past month (van Lee et al. 2012). The questionnaire was expanded to include question that allow for quantitative analyses of total energy consumption (in kilojoules) and macronutrients. The macronutrients were used as measures of quantitative dietary intake.

**Table 1:** Demographic data of study participants.

| Characteristic | Value |
| --- | --- |
| Age (years; mean ± 1 SD) | 9.34 ± 0.88 |
| Gender (F/M) | 13 / 18 |
| BMI (kg/m<sup>2</sup>; mean ± 1 SD) | 16.93 ± 1.98 |

SD: standard deviation

### 2.2 Assessment of executive functioning

Participants were invited to the Donders Institute for Brain, Cognition, and Behaviour (Nijmegen, the Netherlands) together with one of their parents for three visits with additional assessments (reported elsewhere). This study used the first visit (intake session) for collection of exhaled breath sample and executive functioning measures. During this visit, consent forms were signed. Children first completed a general fluid intelligence task, followed by the Eriksen flanker task (Eriksen and Eriksen 1974). In this child-adapted task, participants were presented with a string of five fish on a computer screen and were asked to respond to the orientation of the middle fish by pressing the left or right arrow on the keyboard, correspondingly. The flanking fish could either be oriented in the same direction (congruent trials) as the middle fish, or in the opposite direction (incongruent trials) of the middle fish. The task included 40 incongruent and 40 congruent stimuli, randomly distributed and with an equal amount of left and right responses for each condition.

The flanker error rate and flanker reaction time measures were used to assess executive functioning. The flanker error rate refers to the number of errors made, i.e. selecting the wrong direction of the target, when it was flanked by distractors pointing in another direction (incongruent) versus targets flanked by stimuli pointing in the same direction (congruent). The flanker reaction time is the average time of a child to select the target in the incongruent condition versus the congruent condition. Next, the compatibility effect based on reaction time was calculated by subtracting the average of the correct congruent trials from the average reaction time of the correct incongruent trials. To calculate the compatibility effect based on error rate, false and correct answers were respectively scored with 1 and 0. The average was taken for all the congruent trials as well as the incongruent trials. The error rate compatibility effect was calculated by subtracting the error rate of the congruent trials from the error rate of the incongruent trials. A higher compatibility effect in either errors or reaction times indicates more difficulty with the task (incongruent trials in particular) and lower executive functioning abilities.

### 2.3 Breath sampling

After performing the Eriksen flanker task, end-tidal breath samples were collected using a Loccioni® breath sampler (Angeli di Rosora, Italy) as shown in 2. Children practiced exhaling into an open-end heated calibrated buffer pipe equipped with a mouthpiece with bacterial filter (GVS, Morecambe, UK) and a CO_2_ sensor (CAPNOSTAT5, Hamilton Medical, Zeewolde, the Netherlands). Each child provided 2-3 exhalations at a controlled flow rate of 50 mL/s that was maintained via a colour scale visualisation on the screen of the breath sampler. A discard bag (150 mL) was included to allow collection of the end-tidal part of the breath in duplicate into a 3-L Tedlar® sample bag. Two additional 3-L Tedlar® bags were filled with air from the sampling room to use as background reference.

### 2.4 Analysis of exhaled breath VOCs

#### 2.4.1 Untargeted exhaled breath profiling using Proton Transfer Reaction – Mass Spectrometry

Untargeted breath profiling was carried out on an in-house developed Proton Transfer Reaction-Mass Spectrometry (PTR-MS) instrument. Details of the PTR-MS instrument are described elsewhere (Boamfa et al. 2004; Steeghs et al. 2006). Tedlar® bags were connected to a 1/8” heated PFA inlet line (60°C), which sampled the gaseous samples into the drift tube of the PTR-MS at a rate of 0.6 L/h. The pressure, temperature, and voltage in the drift tube were set to 2.05 mbar, 60°C, and 550 V, respectively resulting in a reduced electric field (*E/N*) of approximately 131 Td. The scan range was set to *m/z* 20-160. The dwell time was set at 1 s resulting in a scan rate of 1 amu/s. Data was acquired for three full scan range cycles, which were subsequently averaged.

Tedlar® bags filled with room air bags were used to determine the background level for each ion. Analysis of room air bags was done in the same manner as for breath samples. The limit of detection (LOD) was determined using the definition of the International Union of Pure and Applied Chemistry (IUPAC) guidelines (McNaught and Wilkinson 1997). Briefly, the LOD of an ion is the average background level (room air) plus three standard deviations. Annotation of ions was done based on *m/z* values and literature (Pagonis et al. 2019). All ion counts were normalised to the reagent ion isotopologue (H_3_^18^O^+^; *m/z* 21; ×500) signal after which normalised counts were converted into part-per-billion volume (ppbV) concentration values as outlined by De Gouw *et al*. (de Gouw et al. 2003).

#### 2.4.2 Analysis of breath ethylene and methane

Ethylene measurements were performed on a laser-based photoacoustic detector (ETD-300, Sensor Sense B.V., Nijmegen, the Netherlands). The ETD-300 has been used before for breath analysis (Cristescu et al. 2014; Paardekooper et al. 2017). Tedlar® bags with breath samples were connected to an in-house designed sampling system consisting of a pump and a Flow-IQ mass flow controller (Bronkhorst, Ruurlo, the Netherlands) allowing to introduce the breath samples into the ETD-300 at a flow rate of 1.0 L/h. Prior to entering the inlet of the spectrometer, CO_2_ and water vapour were removed using soda lime (Fisher Scientific, Bleiswijk, the Netherlands) and anhydrous calcium chloride (Sigma-Aldrich, Amsterdam, the Netherlands), respectively. Breath samples were analysed for 5 minutes, the average concentration was calculated over the last 2-minute interval.

Exhaled methane was analysed using continuous-wave optical parametric oscillator (cw-OPO)-based photoacoustic spectroscopy (power 1.2 W, tunability 3–4 µm). Technical details of the setup are described elsewhere (Arslanov et al. 2011). Tedlar® bags with breath samples were connected to a pump through 1/8’’ perfluoroalkoxy alkane (PFA) tubing. The flow rate of the pump was set to 1 L/h by using a mass flow controller (Brooks Instruments, Veenendaal, the Netherlands). Data was acquired at a laser scanning rate of 1 kHz and an integration time of 0.25 s.

### 2.5 Statistical analysis

All exhaled breath VOC, FFQ, and Flanker task results were combined in one master file. For both the flanker error rate and the flanker reaction time, a paired *t*-test was used to examine if there was a significant difference in means for the congruent trials and incongruent trials. To assess the predictive value of breath profiles and macronutrient intake on Flanker task output, partial least square (PLS) regression was used to determine significant features. Prior to PLS regression, missing value were replaced using mean value imputation (Wei et al. 2018) and subsequently *zscore* normalised. The optimal number of latent variables (LVs) was selected based on the lowest mean square error (MSE) of the prediction using leave-one-out cross validation (Godlewski et al. 2023). Variable selection was done using the significant multivariate correlation (sMC) metric at an α of 0.05 (T. N. Tran et al. 2014). Regression coefficients (β) were used to explore potential explanations for the relation between breath VOCs, macronutrient intake and Flanker task performance. PLS models for the error rate and reaction time compatibility effects were validated using permutations.(Westerhuis et al. 2008)

## 3. Results & Discussion

### 3.1 Executive functioning

Twenty-six children completed the flanker task, including congruent and incongruent trials. Children made fewer errors on average during congruent trials than during incongruent trials (p < 0.001; Figure *3*A). Similarly, children responded faster to congruent trials than to incongruent trials (p < 0.001; Figure *3*B). These findings align with prior research, demonstrating that accuracy and response time are better in congruent trials (Bulger et al. 2021; Chiu et al. 2017; Stins et al. 2007). There was no difference in flanker compatibility effects between boys and girls (error rate: p = 0.7319; reaction time: p = 0.7947). In addition, age was not found to be a significant factor for either the flanker compatibility effect in error rates (p = 0.1419) or in reaction time (p = 0.2015). No correlation was found between the compatibility effects in reaction time and error rate (p = 0.9190), hence, both measures were considered as two separate indicators of (decreased) executive functioning.

**Figure 1:**
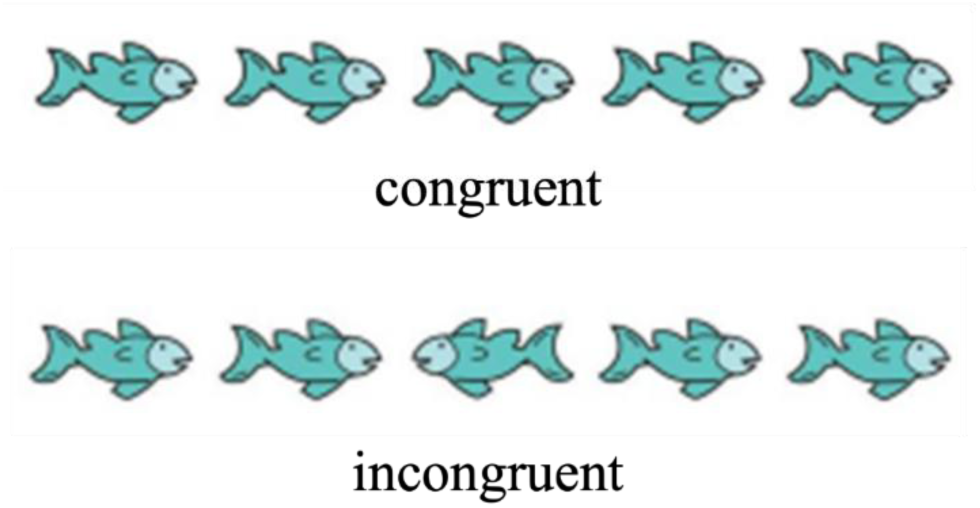
An adapted version of the Eriksen flanker task. Children were instructed to respond to the orientation of the middle fish by pressing a left or right button. The middle fish could point in the same direction as the flanking fish (congruent condition) or in the opposite direction (incongruent condition). The flanker compatibility effect was calculated by subtracting congruent response times or error rates from the incongruent ones, with a larger compatibility effect indicating more difficulties with executive functioning.

**Figure 2:**
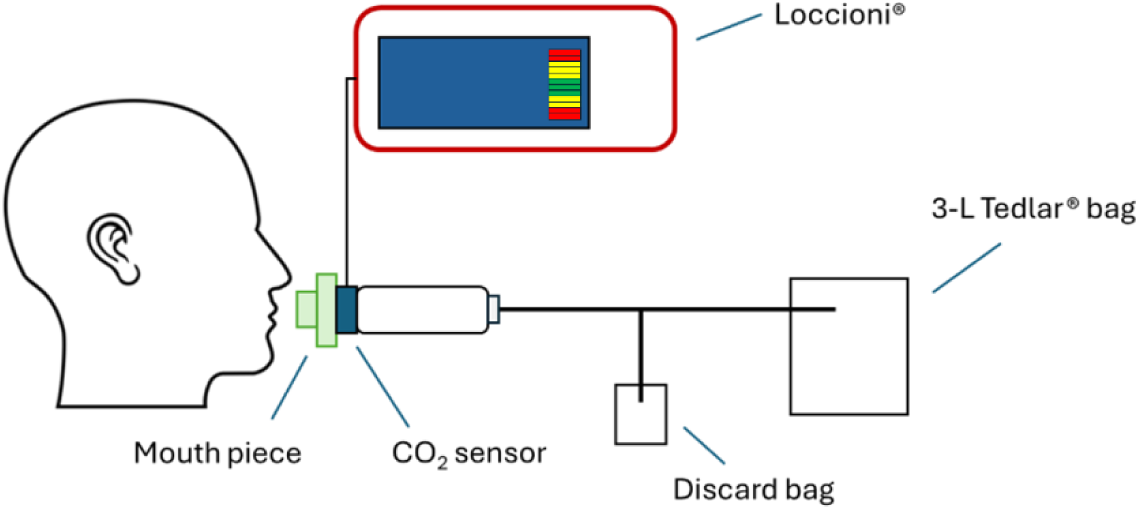
Schematic representation of the setup used for exhaled breath collection.

**Figure 3:**
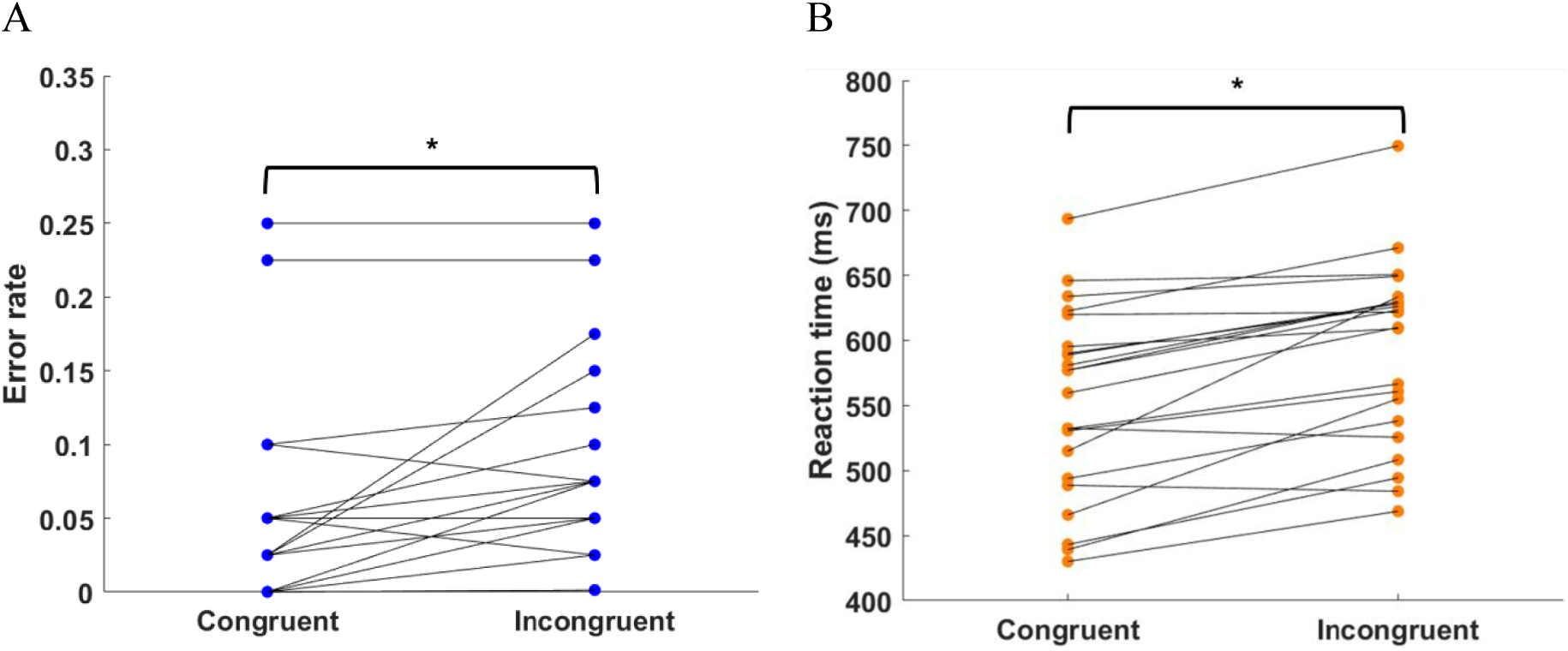
Performance of children during congruent and incongruent trials of the Flanker task. (A) Error rate; (B) Reaction time. *: p < 0.05

### 3.2 Exhaled breath volatiles as non-invasive indicators for executive functioning

Exhaled breath VOCs (Table *2*) and FFQ data (Table *3*) were used as predictor variables to assess associations with the compatibility effects of the flanker test. As ions can only be assigned a putative identity based on their *m/z* value, literature on breath research was used to aid in the assignments. The aim here is to explore whether breath metabolites reflect processes that could affect cognitive functioning. To achieve this, two separate PLS regression models were generated: one for the compatibility effect in error rates and another for reaction time. PLS models were found to be statistically significant through permutation testing (error rate: p = 0.020; reaction time: p = 0.008). In total, 5 and 3 out of 17 included variables in the PLS model (breath VOCs and macronutrient intake) were found to be significantly associated with performance in the flanker test (sMC: p < 0.05) for the compatibility effect in reaction (Figure *4*A) and error rate (Figure *4*C), respectively. The effect of the variables on the compatibility effect is given by the regression coefficient (Figure *4*B-D). A positive regression coefficient refers to an increase of the compatibility effect, i.e. a reduction in executive functioning, whilst a variable with a negative regression coefficient is associated with an improvement in executive functioning.

**Figure 4:**
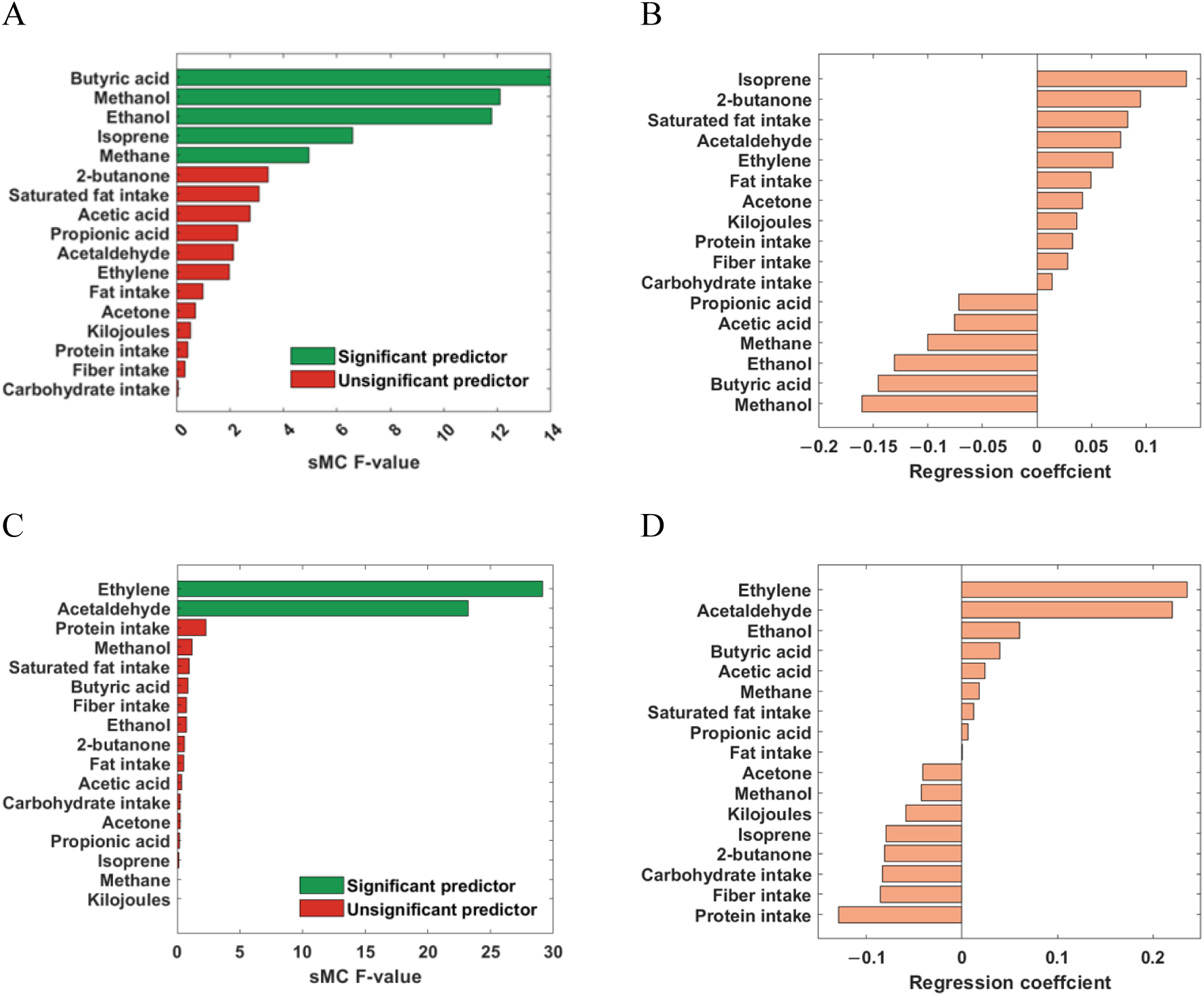
Results of PLS regression. (A) Variable selection using sMC for the compatibility effect in reaction time, and (B) the corresponding regression coefficients for the reaction time model. (C) Variable selection using sMC for the compatibility effect in error rates, and (D) the corresponding regression coefficients for the error rate model.

**Table 2:**
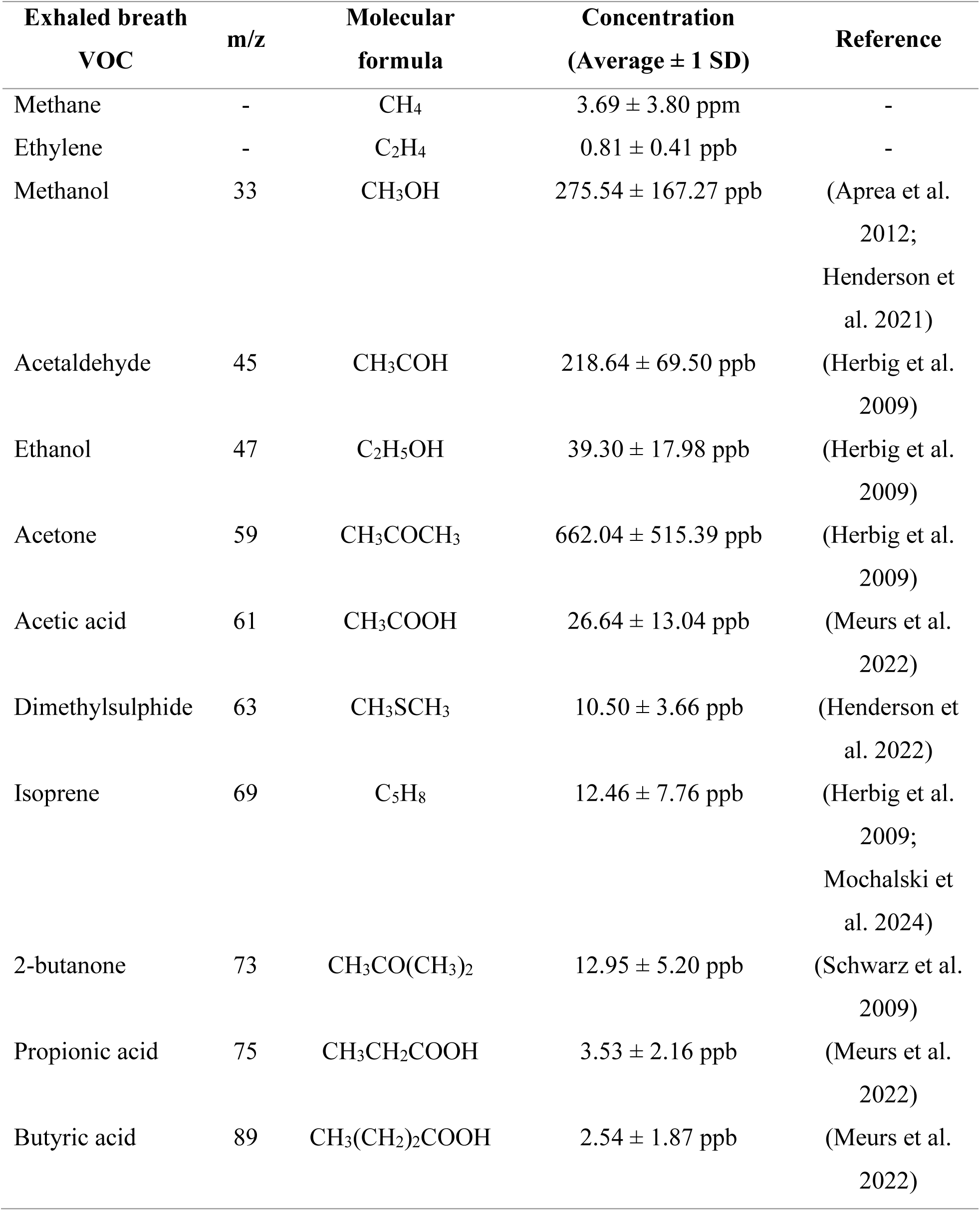
Overview of measured breath VOCs and their descriptive statistics.

| Exhaled breath<br>VOC | m/z | Molecular<br>formula | Concentration<br>(Average ± 1 SD) | Reference |
| --- | --- | --- | --- | --- |
| Methane | - | CH<sub>4</sub> | 3.69 ± 3.80 ppm | - |
| Ethylene | - | C<sub>2</sub>H<sub>4</sub> | 0.81 ± 0.41 ppb | - |
| Methanol | 33 | CH<sub>3</sub>OH | 275.54 ± 167.27 ppb | (Aprea et al.<br>2012;<br>Henderson et<br>al. 2021) |
| Acetaldehyde | 45 | CH<sub>3</sub>COH | 218.64 ± 69.50 ppb | (Herbig et al.<br>2009) |
| Ethanol | 47 | C<sub>2</sub>H<sub>5</sub>OH | 39.30 ± 17.98 ppb | (Herbig et al.<br>2009) |
| Acetone | 59 | CH<sub>3</sub>COCH<sub>3</sub> | 662.04 ± 515.39 ppb | (Herbig et al.<br>2009) |
| Acetic acid | 61 | CH<sub>3</sub>COOH | 26.64 ± 13.04 ppb | (Meurs et al.<br>2022) |
| Dimethylsulphide | 63 | CH<sub>3</sub>SCH<sub>3</sub> | 10.50 ± 3.66 ppb | (Henderson et<br>al. 2022) |
| Isoprene | 69 | C<sub>5</sub>H<sub>8</sub> | 12.46 ± 7.76 ppb | (Herbig et al.<br>2009;<br>Mochalski et<br>al. 2024) |
| 2-butanone | 73 | CH<sub>3</sub>CO(CH<sub>3</sub>)<sub>2</sub> | 12.95 ± 5.20 ppb | (Schwarz et al.<br>2009) |
| Propionic acid | 75 | CH<sub>3</sub>CH<sub>2</sub>COOH | 3.53 ± 2.16 ppb | (Meurs et al.<br>2022) |
| Butyric acid | 89 | CH<sub>3</sub>(CH<sub>2</sub>)<sub>2</sub>COOH | 2.54 ± 1.87 ppb | (Meurs et al.<br>2022) |

**Table 3:**
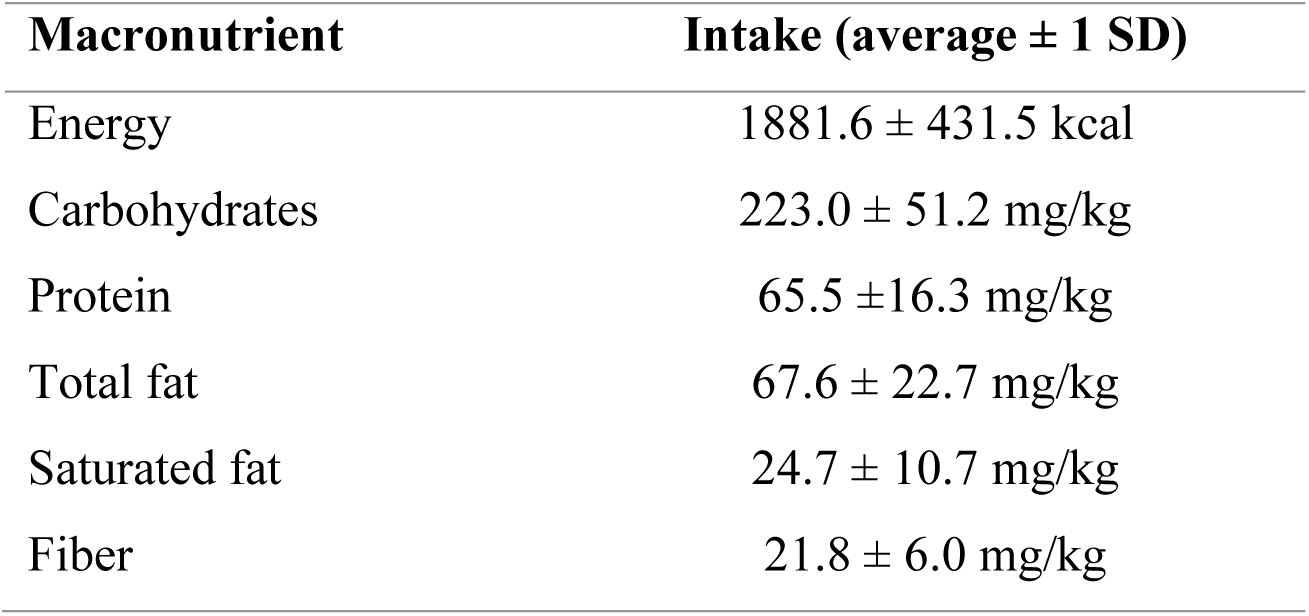
Descriptive statistics for macronutrients.

To gain additional insights into the association between breath VOCs and executive functioning, the biological functions per breath VOC were retrieved. This information is listed in Table 4 along with their association with executive functioning.

**Table 4:** VOCs with statistically significant association to executing functioning performance and their physiological role in relation to cognitive function or gut microbial activity.

| VOC | Physiological origin | Association with executive functioning<sup>a</sup> | References |
| --- | --- | --- | --- |
| Acetaldehyde | Ethanol metabolism, lipid peroxidation | - | (Tacconi et al. 2017) |
| Butyric acid | Fermentation of dietary fibre and non-digestable polysaccharides | + | (Silva et al. 2020) |
| Ethanol | Macronutrient metabolism by the gut microbiome | + | (Oliphant and Allen-Vercoe 2019) |
| Ethylene | Lipid peroxidation | - | (Paardekooper et al. 2017) |
| Isoprene | Cholesterol biosynthesis via the mevalonate pathway within the liver and muscular lipolytic cholesterol metabolism | - | (Deneris et al. 1984; Mochalski et al. 2024) |
| Methane | Anaerobic fermentation of carbohydrates by enteric microflora | + | (Dorokhov et al. 2015) |
| Methanol | Pectin degradation, demethylation of endogenous cellular proteins for regulation, and vitamin B12 synthesis | + | (Dorokhov et al. 2015) |

<sup>a</sup>: Association of VOC with executive functioning, with ‘+’ meaning better (i.e., smaller Flanker effect) and ‘-’ meaning worse (i.e. larger Flanker effect) executive functioning with an increase in the VOC, respectively.

Several VOCs present in breath samples can be related to gut microbial activity as they are known to be produced by the gut microbiota, and it is reasonable to assume that they can be exhaled in breath after absorption from the colon, systemic circulation in blood and then excretion in the lungs (Smolinska et al. 2018). Butyric acid is one of the three most abundant SCFAs which is formed through the fermentation of non-digestible polysaccharides and dietary fibre by the gut microbiota (Ahmed et al. 2022). Whilst the main route for excretion of SCFAs is via the lungs after oxidation to CO_2_ (Boets et al. 2017), prior studies have shown the possibility of detecting butyric acid, among other SCFAs, in exhaled breath (Meurs et al. 2022; Neyrinck et al. 2021). Butyric acid is a metabolite of significant interest as it has been reported for its advantageous effects on gut homeostasis and energy metabolism (Liu et al. 2018). Previous research has suggested that this anti-inflammatory effect can be linked to improved executive functioning (Moriguchi and Hiraki 2013). Moreover, a study in mice found that butyrate intake reversed the detrimental effects of a low-fibre maternal diet on cognitive functioning in the offspring (Yu et al. 2020). This is supported by the current finding of a negative regression coefficient (i.e. reducing the reaction time compatibility effect) in the PLS model, indicating better executive functioning associated with breath butyrate levels. The impact of SCFAs on executive functioning has also been described in studies related to Parkinson’s disease (PD) since one of the PD symptoms is the loss of executive functioning (Sasso et al. 2023). Interestingly, the abundance is reduced of SCFA-producing *Prevotellaceace* in the colon in faecal samples of PD patients (Scheperjans et al. 2015). In addition, butyrate-producing species *Blautia*, *Coprococcus*, and *Roseburia* were also found to be less abundant in faecal matter of PD patients (Keshavarzian et al. 2015; Spielman et al. 2018)

Acetic acid and propionic acid were also measured in the exhaled breath of the children and displayed the same association with the reaction time as butyric acid. This supports the potential positive association between SCFAs and executive functioning. Increased consumption of dietary fibre can lead to higher SCFA production by the gut microbiota (Sasso et al. 2023), a correlation between children’s fibre intake and exhaled concentrations of SCFAs was expected. A positive correlation was indeed observed, however, the correlation was not significant (Spearman’s r: 0.3250; p = 0.0749).

In this study, methane (CH_4_) concentration in breath samples was also significantly associated with improved executive functioning. In the gut, CH_4_ is mainly produced by *Methanobrevibacter smithii* and *Methanosphaera stadtmanae* through reduction of formate (HCO_2_-) and CO_2_, respectively (Smith et al. 2019). *Methanobacterium ruminatum* and certain species of *Clostridium* and *Bacteroidetes* produce CH_4_ as well (Roccarina et al. 2010). Previously, lower abundance of numerous *Bacteroidetes* in the gut microbial community has been associated with reduced inhibitory control (a core executive function) in a mouse model (Arnoriaga-Rodríguez et al. 2021). Together with the current findings, this might indicate that circulating CH_4_ levels could impact executive functioning in humans. However, more research will be needed to further elucidate a potential association between CH_4_ and inhibitory control.

We also observed a positive association between both ethanol and methanol levels in breath samples and executive functioning (i.e., reduced Flanker effect). This finding aligns with previous research that linked several bacterial strains known to produce methanol and/or ethanol to executive functioning. Endogenous ethanol and methanol both originate from carbohydrate fermentation in the colon (Dorokhov et al. 2015; Oliphant and Allen-Vercoe 2019). It has been demonstrated that different strains of *Clostridium* produce both methanol and ethanol, among other volatile products (Kuppusami et al. 2015; Stotzky et al. 1976). Cox *et al*.(Cox et al. 2021) found an unclassified *Clostridium* strain, among other bacteria, to be positively correlated with cognitive functioning in multiple sclerosis patients. Additionally, the genera mentioned above in relation to PD, namely *Blautia*, *Coprococcus*, and *Roseburia* also produce ethanol (Oliphant and Allen-Vercoe 2019).

Breath isoprene has been related to muscular lipolytic cholesterol metabolism and cholesterol metabolism via the mevalonic acid pathway (Deneris et al. 1984; Mochalski et al. 2024). To the best of our knowledge, there has been no report on the production of isoprene by the gut microbiome. Exploring the microbial VOC (mVOC) database (Lemfack et al. 2018) for isoprene also did not reveal any hits for gut microbial species. Increased levels of exhaled isoprene were also observed as a response to stress during a cognitive test. It is thought that cortisol production is related isoprene levels, however, the exact pathway is not known (Gall et al. 2021). That observation is in line with the present study as higher isoprene concentrations in breath were associated with increased reaction time compatibility effect according to the PLS model. Possibly, children who experienced more difficulty during the flanker task also experienced more psychological stress, which was reflected in higher levels of breath isoprene right after the flanker test when the breath samples were collected.

Both acetaldehyde and ethylene have been reported as (by-)products of inflammatory processes (Moeskops et al. 2006; Paardekooper et al. 2017). Acetaldehyde is also formed as an intermediate product of ethanol and pyruvate metabolism and has been linked to DNA damage and inhibition of fatty acid oxidation (Comporti et al. 2010; Tacconi et al. 2017). Ethylene is a marker of lipid peroxidation and can be induced by the release of reactive oxygen species triggered by an immune response (Paardekooper et al. 2017). In the present study, increased levels of both acetaldehyde and ethylene in breath were related to an enhanced compatibility effect, indicating worse executive functioning. Indeed, inflammatory processes have been found to impact neurocognitive processes during similar executive functioning tasks, such as the Stroop task, in adults (Brydon et al. 2008). Additional evidence for a potential link between breath ethylene and reduced executive functioning can also be found in literature in relation to PD. Parkinson’s patients showed an increased abundance of *Enterobacteriaceae* species, contributing to higher serum levels of lipopolysaccharide (LPS) (Roy Sarkar and Banerjee 2019; Scheperjans et al. 2015). A study by Paardekooper *et al*.(Paardekooper et al. 2017) demonstrated the formation of ethylene as a result of intravenous LPS administration. Higher levels of exhaled ethylene in this study could therefore indicate LPS-induced inflammatory processes, relating to a negative effect on executive functioning. Also, acetaldehyde can be linked as evidence to stress response. Tsuruya *et al*.(Tsuruya et al. 2016) indicated *Ruminococcus* to be involved in anaerobic acetaldehyde production from ethanol in the intestine. In a mouse model, following stress induction, among other taxa, *Ruminococccaceae* abundance in the gut microbial community was significantly correlated with anxious behaviour. However, underlying mechanisms linking behaviour and this particular bacterial taxon is not understood yet.

Regarding nutrient intake, from all FFQ measures (i.e. total energy consumption and macronutrient intake) in this study, only protein intake was related to a reduced compatibility effect (in error rates) in children. This is in line with earlier studies in adults where additional intake of protein resulted in improved visuo-spatial short-term memory (K Fischer 2002) and dietary intake of proteins related to better executive functioning (Muth and Park 2021). For the compatibility effect in reaction times, none of the macronutrients was found to play a significant role. In other studies, fat, protein, and carbohydrate consumption also showed neither a positive nor a negative effect on choice reaction time (Karina Fischer et al. 2001). However, a systematic review suggests associations between dietary (macronutrient) consumption and executive functioning in children, with e.g. negative relations found between (saturated) fat intake and executive functioning (Cohen et al. 2016). Yet, another study in adolescents found little evidence for associations of macronutrient composition with attention capacity (Henriksson et al. 2017). The more objective measurement of volatiles might lead to more consistent results in the future than subjective, parent-reported food intake, as in the FFQ (Kristal et al. 2005).

As this is a cross-sectional study, individual VOCs and nutrient intake cannot be directly and causally linked to a change in cognitive function. Neither did the current study include a profiling of the gut microbiota and related metabolites in faeces or blood. The gut microbiome is thought to play an important role function across the GBA (Zhang et al. 2020). Therefore, in this exploration between breath VOCs and executive functioning, the main focus was around metabolite production by the gut microbiome. However, it should be noted that volatile metabolites can originate from other sources in the body and concentration are affected by various external factors as lifestyle and environmental exposure (Blanchet et al. 2017). In addition, it should be noted that the breath VOCs are not exclusively produced within the gut, but can originate from other sources within the body. To further investigate relations between VOCs, food intake and executive cognitive functioning, objective recordings and longitudinal or intervention designs need to be considered. Furthermore, the concentration of some gut microbiome-related VOCs, such as CH_4_ and SCFAs, might vary in the period after the last meal intake (Neyrinck et al. 2021). Therefore, the time between the last meal intake and exhaled breath collection should be standardised.

## 4. Conclusion

In conclusion, exhaled breath analysis bears potential for studying associations between gut microbiota, as well as host inflammatory pathways and cognitive functioning in children. This opens new endeavours by offering a non-invasive tool for further investigation of gut-brain health-related topics. Elucidating biochemical changes reflected in exhaled breath can provide insights for developing novel, strategies to positively impact cognitive function.

## Acknowledgements

This work has been supported by the European Union’s Horizon 2020 Research and Innovation Program through the NUTRISHIELD project (https://nutrishield-project.eu/) under Grant Agreement No. 818110; The European Union’s EFRO grant, case number PROJ-00405; and by Reckitt/Mead Johnson Nutrition, Nijmegen, The Netherlands.

## Author contributions

**JM:** Methodology, Data Curation, Formal Analysis; Writing (Original Draft). **BH:** Methodology, Data Curation; Formal Analysis **CvD:** Methodology, Data Curation; Formal Analysis. **GLB:** Data Curation; Formal Analysis. **ES:** Data Curation, Writing (Review & Editing). **JAD:** Supervision; Writing (Review & Editing). **GG:** Supervision; Writing (Review & Editing). **EA:** Funding acquisition; Conceptualisation; Supervision; Writing (Review & Editing). **SMC:** Funding acquisition; Conceptualisation; Supervision; Writing (Review & Editing).

## Data availability

The data is available upon request to the corresponding author

